# Proteomic clocks reveal organ-specific signatures of aging in mice

**DOI:** 10.64898/2026.10.27.762011

**Authors:** Sean Lim, Andrew M. Pickering

**Affiliations:** Department of Biosciences, Rice University, Houston, TX, 77251, USA; Department of Integrative Biology and Pharmacology, The University of Texas Health Science Center at Houston, Houston, TX, 77030, USA; Institute on Aging, The University of Texas Health Science Center at Houston, Houston, TX, 77030, USA

**Keywords:** aging clock, proteomics, organ aging, transcriptome, spatial transcriptomics, mouse

## Abstract

Aging alters protein abundance throughout the body, but whether proteomic aging signatures are shared across organs, and how they are represented in RNA, remains unclear. Plasma proteomics does not measure protein abundance directly within individual organs. Here, we developed organ-specific proteomic aging clocks for eight mouse organs. The clocks retained age associations across independent cohorts but transferred poorly between organs of the same mice, indicating predominantly organ-specific aging signatures. This organ specificity extended to RNA. Applied to the transcriptomes of the genes encoding each clock’s protein features, with protein-derived weights unchanged, the scores retained age associations in independent transcriptomic cohorts, with stronger associations in the corresponding organs. The strength of age associations across molecular layers also differed by organ: in paired analysis, protein scores tracked age more strongly than RNA scores in brain and skeletal muscle, with no statistically detectable difference in kidney, liver or lung, suggesting a greater post-transcriptional contribution to proteomic aging in brain and muscle. In spatial transcriptomes, protein-derived RNA scores rose with age within tissue niches, usually without detectable changes in relative niche abundance. In the brain, white matter and glia-enriched microdomains showed elevated proteomic scores compared to other regions since young age. Together, these findings support organ-specific regulation of proteomic aging and identify molecular remodeling within persistent tissue compartments.

## 1. Introduction

Aging changes molecular composition throughout the body, but organs do not age at the same rate^1,2^. The age of a single organ can therefore carry information that whole-body measures miss. Organ ages estimated from human plasma proteins, for example, are associated with future disease in the corresponding organ^1^. Most such estimates come from molecular aging clocks, which are models that combine many age-associated molecular features into a single score that predicts age. DNA methylation clocks are the most developed class; they predict chronological ages tissues in several species^3,4^ and have been shown to respond to lifespan-extending interventions^5^. Transcriptomic clocks use gene expression instead^6,7^, and proteomic clocks, a more recent approach, use protein abundance to estimate age^8^. In human plasma, proteomic clocks have been linked to incident disease and mortality^9^.

Most existing aging clocks have been developed using plasma, largely because blood collection is minimally invasive, permits repeated measurements in living individuals, and is readily applicable to large clinical cohorts. However, plasma-based organ clocks, can only infer organ aging indirectly from circulating proteins assigned to organs by tissue-expression annotations^1^, whereas tissue proteomics allows clocks to be trained instead on proteins measured in the organ itself^2^. Mice are well suited to this approach, because many organs can be collected from the same animals at defined ages, and independent cohorts and lifespan-extending interventions are available for testing. Age has been predicted from mouse liver proteins^10^, and multi-organ atlases now document age-associated protein changes across many tissues^11–14^. These atlases also report changes shared between organs, notably in immunity and metabolism^12,13^. It is not known whether such shared changes lead clocks from different organs to share systemic proteomic aging trajectory. If they do, a clock trained in one organ might read age in another; if not, each organ needs its own clock.

Protein abundance is not set by RNA alone, because translation and turnover also determine how much protein a cell contains^15^. Paired measurements in mice and humans have shown that the two layers (transcriptomes and proteomes) diverge further with age^2,11^, but these comparisons were made largely protein by protein. How much of a proteomic aging signature, which integrates many proteins, is also present in RNA, and whether this differs between organs, remains unclear. Comparing a proteomic clock with a transcriptomic clock cannot answer this, because separately trained models select different features, and any difference between them may come from the models rather than the molecular layers. Tissue-trained proteomic clocks allow a more direct test. Each clock is a fixed list of genes with weights, so it can be applied unchanged to the RNA of the same genes in the same organ to ask whether the score still follows age, with no features reselected and no model retrained. Within an organ, the protein and RNA scores therefore differ only in what was measured, and how well the signature transfer indicates how much of the age-related change in that organ’s proteome is also present in its transcriptome.

A bulk transcriptomic score, however, cannot show where in a tissue a signature is expressed. It may rise with age because expression changes within existing cell populations, because those populations change in relative abundance, or both, and single-cell aging atlases have documented both kinds of change^16^. Spatial transcriptomics retains the anatomical position of each gene-expression measurement^17^. Spatial aging clocks have revealed region- and cell-type-dependent aging in the mouse brain^18^, and spatial aging atlases now span multiple mouse organs^19^. A protein-derived signature that transfers to RNA could be scored in such data within tissue niches, groups of locations with similar expression profiles. Following both a niche’s score and its share of the tissue across ages would separate a rise within niches from a shift in composition. A comparison across ages would also show whether a region that scores high is changing faster, or simply expresses the clock genes at a higher level from the start. To our knowledge, protein-derived aging signatures have not been examined at this spatial level.

Here, we trained proteomic aging clocks for eight organs of male C57BL/6J mice sampled at six adult ages^12^, and tested them in independent cohorts generated with different mass-spectrometry workflows and in mice given lifespan-extending interventions. We then asked whether the clocks rely on shared or organ-specific proteins, applied their protein-derived weights unchanged to bulk RNA from matching organs, and projected them onto spatial transcriptomes to compare age-related changes in niche scores with changes in niche abundance.

## 2. Results

### 2.1 Organ-specific proteomic clocks track age across independent datasets

To establish organ-specific measures of proteomic age, we trained a separate elastic-net model for each organ using a dataset generated by Scifo and colleagues^12^. It contains proteomes from eight organs (brain, heart, lung, liver, kidney, spleen, skeletal muscle, and testis) collected from 45 male C57BL/6J mice across six age groups: 3 months (n = 7), 5 months (n = 9), 8 months (n = 8), 14 months (n = 9), 20 months (n = 7), and 26 months (n = 5). After data preprocessing, samples from 44 mice per organ were included in the clock analyses (***Fig. 1A***). All eight models predicted chronological age in leave-one-mouse-out cross-validation, with Pearson correlation coefficients ranging from 0.72 in brain to 0.91 in skeletal muscle and mean absolute errors of 2.47–4.26 months (***Fig. 1B***).

**Fig 1.**
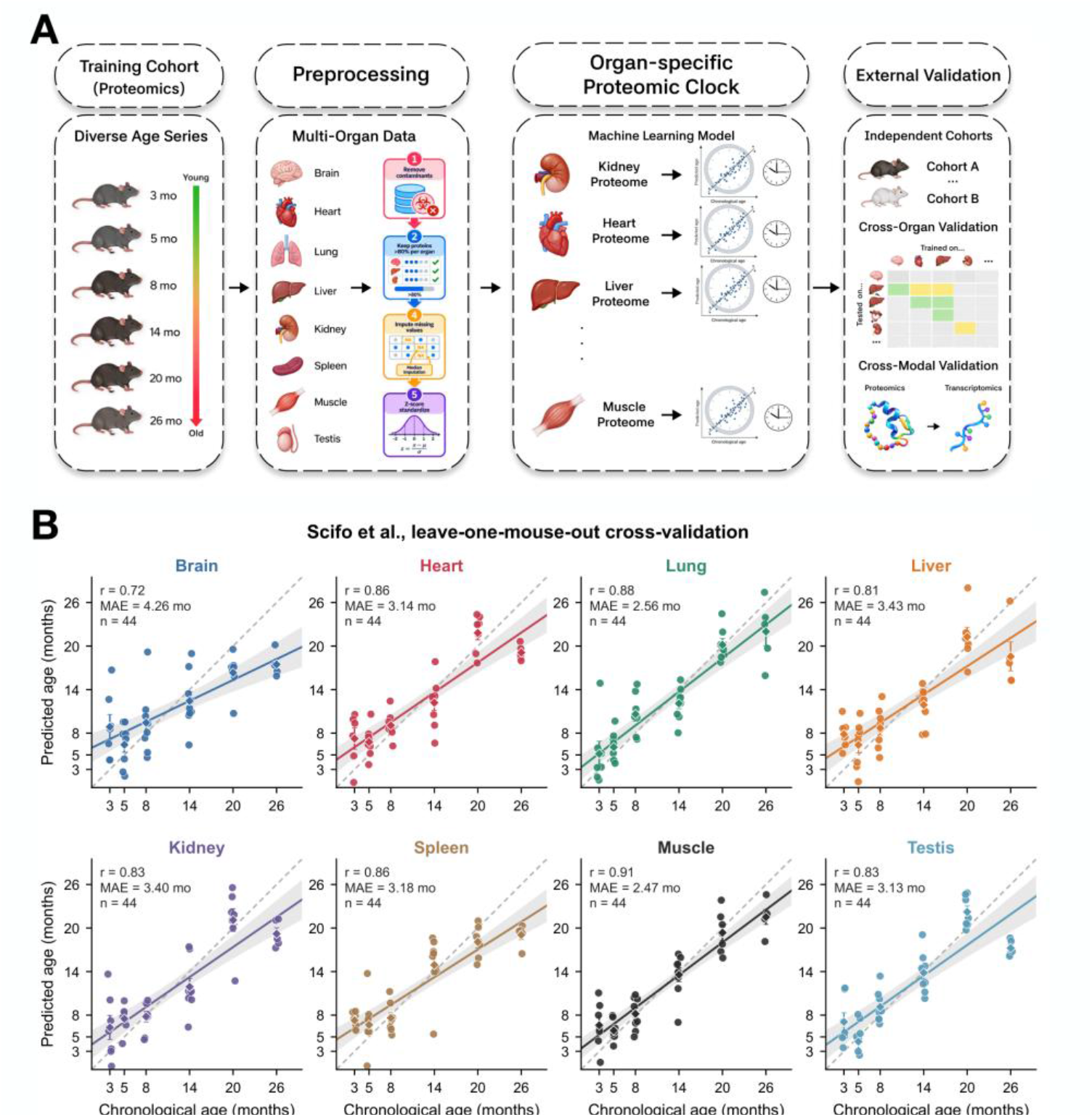
Organ-specific proteomic clocks predict chronological age. **(A)** Study design. Elastic-net clocks were trained on label-free proteomes of eight organs from male C57BL/6J mice aged 3–26 months (Scifo et al.^12^; n = 44 per organ). Fixed core-clock weights were applied without retraining to independent proteomes and RNA; cross-organ proteomic prediction used additional models restricted to proteins retained in all eight organs (Methods). **(B)** Leave-one-mouse-out predicted versus chronological age. Pearson r and mean absolute error (MAE) are shown. Circles, mice; diamonds and error bars, age-group mean ± s.e.m.; lines and bands, linear fit and 95% confidence interval (CI); dashed line, identity.

We next tested whether these age-associated signals transferred to independent datasets generated with different mass-spectrometry workflows. For each organ, we retained proteins selected in more than half of the cross-validation folds as core clock proteins. External mouse datasets were scored using fixed model weights and the standardized abundances of detected core proteins, without refitting. Because protein coverage and abundance scales differed across datasets, we did not convert these scores to ages in months, and asked whether these scores ranked samples by age.

We tested a dataset generated by Takasugi and colleagues^11^, who measured protein abundance in whole tissues of mice aged 6-30 months by tandem mass tag (TMT)-based mass spectrometry. Clock scores increased with age in brain, heart, lung, liver, kidney and skeletal muscle (n = 16 per organ; Spearman rs = 0.81–0.92; all permutation P < 0.001; ***Fig. 2A***), even though only about half to two-thirds of each organ’s core clock proteins were detected (***Supplementary Fig. 1A***). We then used a second independent dataset, from Lu and colleagues^14^, generated with PulseDIA, a label-free approach that improves protein detection by analyzing complementary peptide mass ranges across multiple injections. Here too, scores increased with age in tissues examined (1–24 months; n = 28–31 per tissue; r_s_ = 0.44–0.73; all permutation P ≤ 0.014; ***Fig. 2B; Supplementary Fig. 1B***). This dataset also covered spleen (r_s_ = 0.44) and testis (r_s_ = 0.64), which Takasugi et al. did not profile, and hippocampus, which we scored with the brain clock (r_s_ = 0.55). The clocks therefore retained age-associated information across independent mouse cohorts and distinct quantitative mass-spectrometry methods.

**Fig 2.**
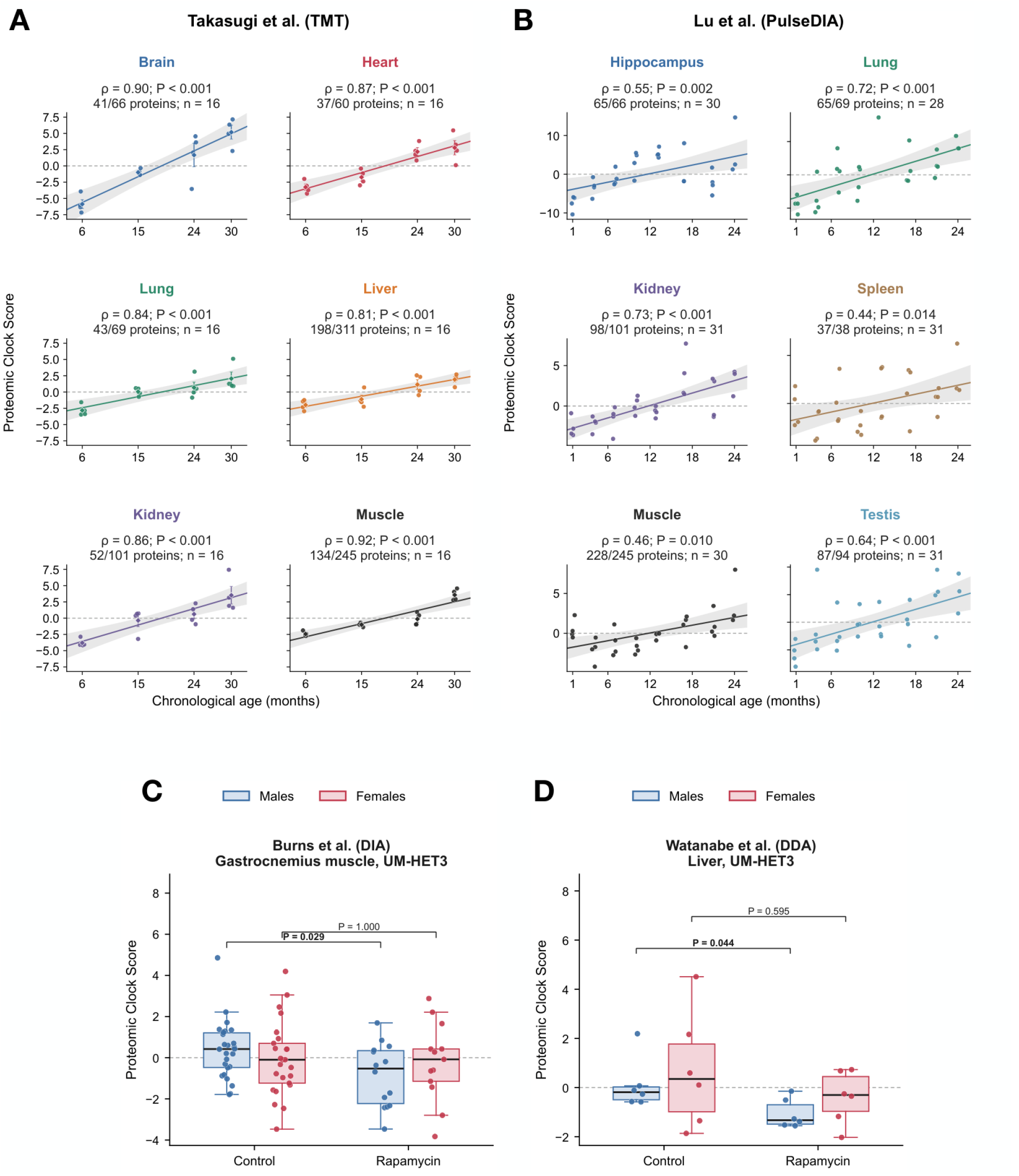
Organ clocks track age in independent proteomes and show variable responses to longevity interventions. **(A)** Clock scores in the Takasugi et al.^11^ TMT cohort (6–30 months; n = 16 per organ). **(B)** Clock scores in the Lu et al.^14^ PulseDIA cohort (1– 24 months; n = 28–31 per tissue); hippocampus was scored with the brain clock. In A and B, annotations give Spearman ρ, two-sided age-permutation P, detected/total core proteins and n; lines and bands show linear fits and 95% CIs. **(C)** Gastrocnemius scores in 12-month-old UM-HET3 mice given rapamycin or control diet (Burns et al.^20^; males, 26 control and 14 treated; females, 25 and 13). **(D)** Liver scores in rapamycin-treated and control UM-HET3 mice (Watanabe et al.^21^; n = 6 per sex and treatment group). In C and D, P values are from two-sided permutation tests of the Mann–Whitney U statistic; bold, unadjusted P < 0.05. Boxes, median and interquartile range; whiskers, 1.5 × IQR. Points represent individual mice. Transferred scores were computed without retraining and are not calibrated ages.

Having established age associations in independent datasets, we next asked whether clock scores also differed under longevity interventions. In a dataset from Burns and colleagues^20^, which included data from UMHET3 mice treated with multiple intervention reported to extended lifespan. We found that rapamycin-treated UM-HET3 mice aged 12 months had lower skeletal-muscle scores than age-matched controls in males (Cliff’s δ = ™0.42; permutation P = 0.029) but not in females (δ = -0.00; P = 1.000), and lower liver scores in females (δ = -0.41; P = 0.038) (***Fig. 2C; Supplementary Fig. 2A***). Rapamycin-treated males also had lower liver scores in the dataset of Watanabe and colleagues^21^ (δ = ™0.72; permutation P = 0.044; n = 6 per group), whereas female scores did not differ significantly from controls (***Fig. 2D***). We also applied the clocks to growth hormone receptor knockout and Snell dwarf mice in the Burns dataset^20^. Growth hormone depletion gave mixed results. In growth hormone receptor knockout (GHRKO) mice, kidney and liver scores tended to be lower than in wild-type controls (δ = ™0.53 and ™0.51; n = 9 knockout and 10 wild-type mice), but neither difference was statistically significant (***Supplementary Fig. 2B***). In Snell dwarf mice, kidney scores were instead higher than in wild-type mice (δ = +0.52; permutation P = 0.047; n = 10 dwarf and 12 wild-type). Clock scores therefore did not respond uniformly to lifespan-extending interventions: rapamycin lowered scores in some organ–sex combinations, but whether and in which direction scores changed varied across datasets and models.

### 2.2 Organ-specific clocks rely on largely distinct protein sets and transfer poorly between organs

To characterize the protein signatures captured by the clocks, we compared core clock proteins with proteins showing significant age-associated abundance changes. Age-differential proteins (Benjamini–Hochberg q < 0.05; ***Supplementary Fig. 3A***) overlapped only partly with the core clock proteins: in spleen, 24 of 38 core proteins were among the 1,792 age-differential proteins, and in kidney, 52 of 101 were among 2,041. The remaining clock proteins did not pass the threshold on their own, yet some contributed substantially to the predictions, including SNF8 and FcRn in liver and ACOT2 in skeletal muscle (***Supplementary Fig. 3B***). The clocks therefore use combinations of proteins, including some without individually significant age associations, rather than only the strongest univariate hits.

**Fig 3.**
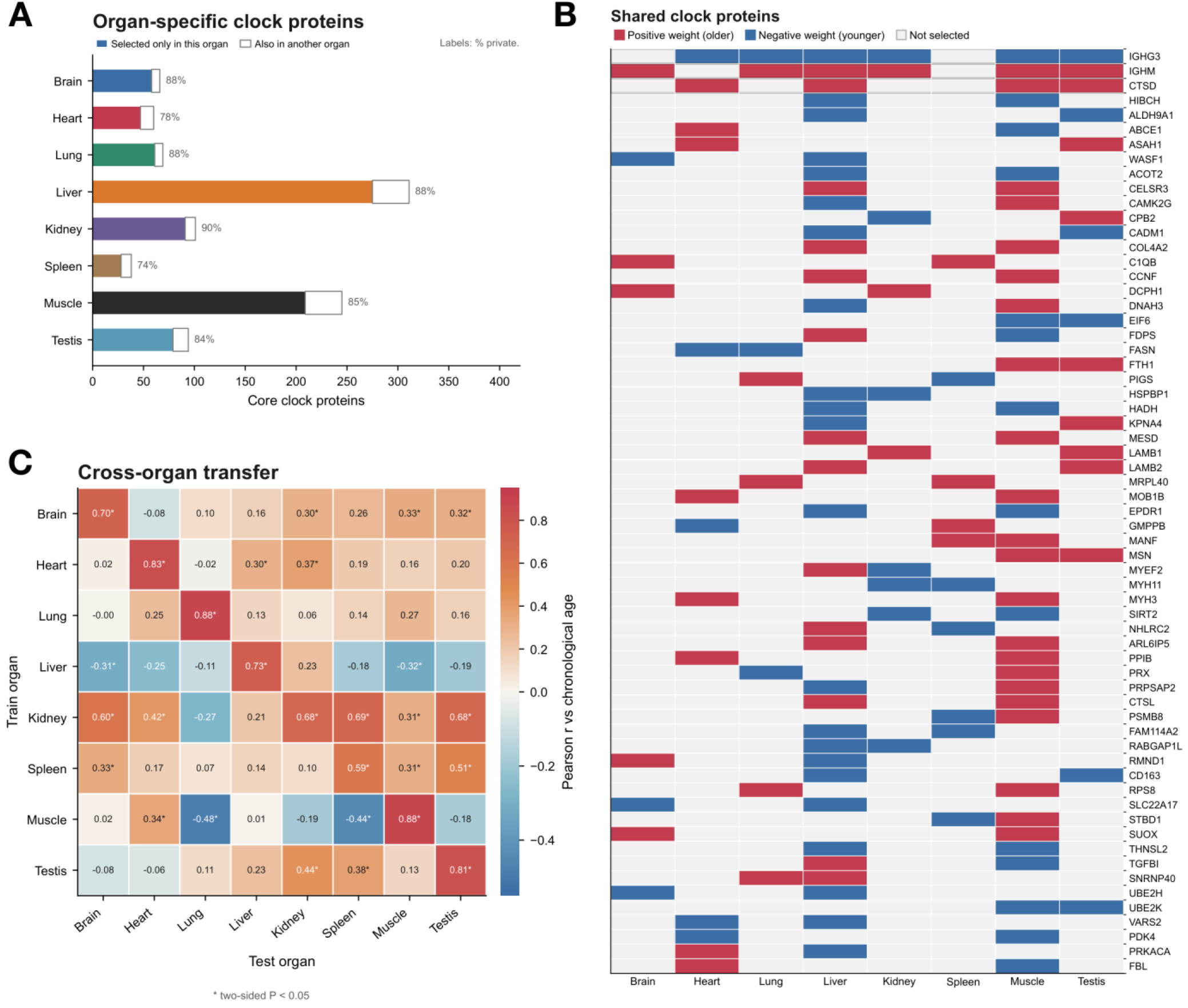
Organ clocks use largely distinct proteins and show limited cross-organ transfer. Analyses used the discovery proteomes of Scifo et al.^12^. **(A)** Core clock proteins selected only in that organ (colored) or also in another organ (white); labels, percentage selected only in that organ. **(B)** Proteins selected in at least two clocks. Red, positive weight (higher abundance, older predicted age); blue, negative weight; grey, not selected. **(C)** Additional models restricted to the 1,074 proteins passing the detection filter in all eight organs were trained in one organ (rows) and applied to another organ of the same held-out mouse (columns). Cells, Pearson r with chronological age (n = 44 mice); diagonal, within-organ performance. *Two-sided P < 0.05, uncorrected.

Across the eight organs, the clocks contained 911 unique core proteins, of which 848 (93%) were selected in just one organ (***Fig. 3A***). Most organ pairs shared 0–5 proteins, the exception being liver and skeletal muscle, which shared 19. Only three proteins were selected in more than two organs: immunoglobulin heavy constant μ (IGHM) and γ3 (IGHG3), each in six organs, and cathepsin D (CTSD), in four (***Fig. 3B***). Their direction was consistent across organs: IGHM and CTSD carried positive weights (higher abundance, older predicted age) wherever they were selected, and IGHG3 carried negative weights.

To test whether the distinct organ-specific protein sets captured a shared aging signature, we trained additional organ-specific models using proteins that passed the detection filter in all eight organs and evaluated their performance in the remaining tissues. Overall, cross-organ transfer was limited, with a median Pearson correlation of r = 0.15 across the 56 organ combinations and only four comparisons showing correlations of r ≥ 0.50 (***Fig. 3C***). Nevertheless, some organ-derived signatures showed broader associations with aging. Most notably, the kidney-derived clock retained relatively strong positive associations with chronological age when applied to brain, spleen, and testis (r = 0.60, 0.69, and 0.68, respectively). The spleen-derived clock also showed a positive association when applied to testis (r = 0.51). In contrast, several cross-organ applications showed weak or even negative associations, including the skeletal-muscle clock applied to lung and spleen (r = ™0.48 and ™0.44, respectively). Transfer was also frequently asymmetric, as illustrated by the kidney-derived clock capturing aging in spleen (r = 0.69), whereas the spleen-derived clock showed little association with kidney aging (r = 0.10). Together, these findings indicate that proteomic aging signatures are predominantly organ-specific, although certain signatures, particularly those derived from kidney, retain age-associated information across multiple tissues.

### 2.3 Protein-derived aging signatures retain age-associated information in bulk RNA

We applied the original gene sets and weights from each organ’s proteomic clock to the corresponding RNA abundances without retraining, testing whether age-associated information learned from proteins was retained at the transcript level.

The dataset generated by Takasugi and colleagues^11^, which we previously used to validate our proteomic clocks, also included transcriptomic measurements from the same mice and organs. Applying our proteomic aging clocks to these RNA data revealed positive associations with chronological age across all six organs examined (n = 16 per organ; Spearman ρ = 0.58–0.92; all permutation P ≤ 0.023; ***Fig. 4A***). The strongest associations were observed in kidney and liver (ρ = 0.92 for both), whereas lung, heart, skeletal muscle, and brain showed weaker, but still significant, correlations (ρ = 0.58–0.78). To determine whether these findings were reproducible, we next applied the same proteomic clocks to an independent transcriptomic dataset generated by Shavlakadze and colleagues^22^, comprising male mice aged 6–27 months. Again, scores increased with age across all five tissues examined (cortex, left ventricle, liver, kidney, and spleen; n = 51–52 per tissue; ρ = 0.63–0.86; all permutation P < 0.001; ***Fig. 4B***). The brain and heart clocks, built from whole-organ proteomes, also worked in cortex and left ventricle, and cortex in fact gave the strongest association (ρ = 0.86). Together, these findings demonstrate that proteomic aging signatures retain age-associated information when applied to RNA across independent cohorts, although the strength of these associations varies considerably between organs and cohorts.

**Fig 4.**
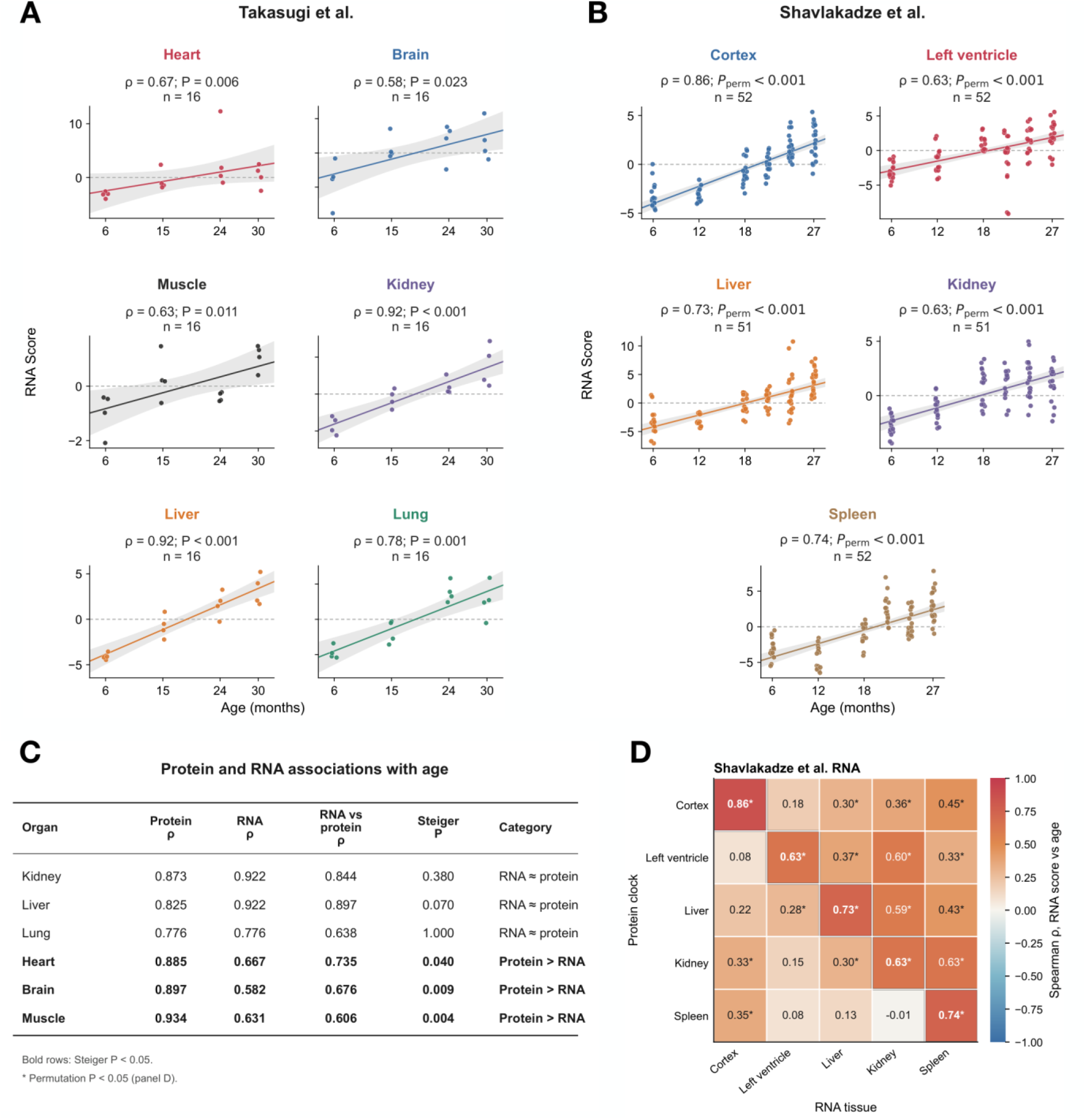
Transfer of proteomic aging signatures to bulk RNA varies by organ. **(A)** Protein-derived RNA scores in the Takasugi et al.^11^ mice shown in Fig. 2A (*n* = 16 per organ), computed from the genes encoding each clock’s proteins with fixed weights. **(B)** The same scores in the Shavlakadze et al.^22^ cohort (6–27 months; *n* = 51–52 per tissue); cortex and left ventricle were scored with the brain and heart clocks. In A and B, annotations give Spearman *ρ* and two-sided age-permutation *P*; lines and bands show linear fits and 95% CIs. **(C)** Protein versus RNA age associations in the Takasugi mice using identical mice, genes and weights; Steiger *P* compares the two correlations (unadjusted; bold, *P* < 0.05). The category “RNA ≈ protein” denotes no statistically detectable difference, not demonstrated equivalence. **(D)** Each clock (rows) applied to RNA from each Shavlakadze tissue (columns); cortex and left-ventricle row labels refer to the brain and heart clocks, respectively. Cells, Spearman *ρ* with age; *two-sided permutation *P* < 0.05, uncorrected. These transferred scores are not calibrated ages.

We tested whether RNA transfer depended on the selected clock genes by comparing the observed age correlations with null distributions generated from random, expression-matched and variance-matched gene sets carrying the same weights (***Supplementary Fig. 4A***). Clock gene sets showed stronger age associations than expected under all three null models in kidney, liver and lung (P ≤ 0.027). Evidence of enrichment over the control gene sets was weaker in heart, brain and skeletal muscle (P = 0.027–0.063).

Paired RNA and protein measurements from the dataset generated by Takasugi and colleagues^11^ allowed direct comparison of age associations using identical mice, genes and clock weights. Protein and RNA scores were correlated in every organ (ρ = 0.61–0.90; ***Fig. 4C***), but they did not track age equally well. In kidney and liver, RNA scores tracked age at least as strongly as protein scores (RNA ρ = 0.92 for both, versus protein ρ = 0.87 and 0.83; Steiger P = 0.38 and 0.070), and in lung the two were identical (ρ = 0.78; P = 1.00). In skeletal muscle, brain, and heart, protein scores showed stronger age associations. Together, these analyses revealed organ-dependent differences in the transcriptional representation of the proteomic aging signatures. No statistically detectable difference in age association was found in kidney, liver or lung, whereas stronger protein associations in brain and skeletal muscle indicate that RNA did not recover the age-associated information in these signatures equally well across organs.

We then asked which features of the clocks carry over to RNA. Keeping only the sign of each weight gave associations close to those of the exact weights in most tissues, although brain and cortex lost some strength (Takasugi brain, ρ = 0.58 to 0.35; Shavlakadze cortex, 0.86 to 0.63; ***Supplementary Fig. 4B***). Giving every gene the same positive weight, which discards sign, weakened the associations in most tissues and reversed them in heart and liver (ρ = ™0.76 and ™0.82 in Takasugi RNA). RNA transfer therefore depended primarily on the selected genes and the signs of their coefficients, rather than their exact magnitudes.

Finally, we asked whether transfer to RNA is organ-specific, as it was for proteins (***Fig. 3C***), by applying each clock to RNA from every other organ (***Fig. 4D***). In the larger Shavlakadze dataset, each clock tracked age best in its own tissue in four of five cases. The exception was the kidney clock, which did equally well in spleen (ρ = 0.63 for both), echoing its transfer to spleen at the protein level. The heart and liver clocks also ranked kidney RNA by age (ρ = 0.60 and 0.59).

### 2.4 Protein-derived aging signatures increase within persistent tissue niches

Bulk RNA shows that the protein-derived signal is present in the transcriptome, but not where in a tissue it resides. We therefore analyzed the 10x Visium aging dataset of Kern and colleagues^19^, which profiled brain, heart, kidney, liver, lung and spleen from male C57BL/6JN mice aged 3.9, 16.8 and 25.8 months, with brain sampled at five coronal levels (bregma ™1.7 to ™4.72 mm). After quality control, the dataset comprised 111 sections from 15 mice, with 10–14 mice per organ and 11 per brain level. Because spots within a section are not independent biological replicates, in every analysis we summed counts across all spots of a section, or of a defined region within it, to obtain one profile per mouse.

We first checked that these data carry an age signal of their own. For each organ and brain level, we trained elastic-net clocks on the 5,000 most variable genes of the summed profiles. In leave-one-mouse-out cross-validation, these organ-specific transcriptomic clocks predicted age in every organ and brain level (Pearson r = 0.61–0.95; MAE = 2.5–5.7 months; all permutation P < 0.05; ***Fig. 5A***). In the brain, error increased steadily from the most anterior to the most posterior level (MAE = 2.5, 2.7, 2.9, 3.6 and 4.6 months for levels 1–5), which may reflect regional differences in transcriptomic aging, although with 11 mice per level this is tentative. Because the dataset has only three ages, we also trained clocks on the 3.9- and 25.8-month mice and predicted the 16.8-month mice. Their mean predicted ages fell between the two training ages in every organ (MAE = 1.4–4.2 months) except lung, where all four middle-aged mice were predicted younger than their true age (6.0–13.1 months; mean, 10.0 months; ***Supplementary Fig. 5***). This model therefore underpredicted age in the held-out middle-aged lung samples. These RNA clocks shared only 0–12 genes per organ with the protein clocks, so they serve as a reference for how much age signal the data contain, independent of the protein-clock genes.

**Fig 5.**
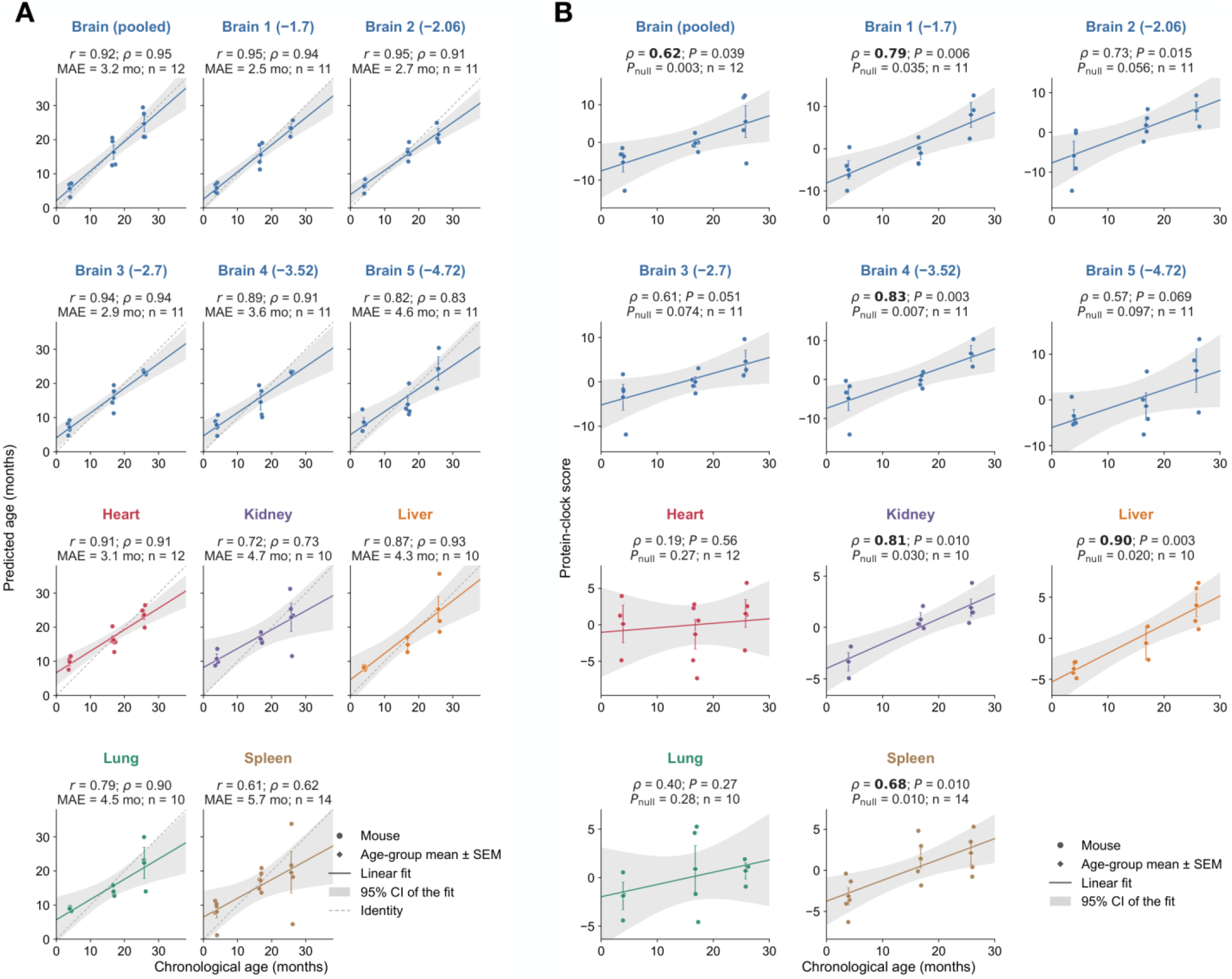
Proteomic aging signatures transfer to spatial transcriptomes in liver, kidney, spleen and selected brain regions. Visium counts were summed per section from male mice aged 3.9, 16.8 and 25.8 months (Kern et al.^19^); brain levels are labelled by bregma (mm), and pooled brain profiles average the available level-centred profiles within each mouse. **(A)** Leave-one-mouse-out predictions of RNA clocks trained on these profiles. **(B)** Protein-derived RNA scores computed from the same profiles with fixed proteomic weights and without retraining. In B, *P* denotes the two-sided age-permutation test and *P*null the one-sided comparison with 1,000 expression-matched gene sets; bold *ρ* indicates *P* < 0.05 in both tests. Circles, mice; diamonds and error bars, age-group mean ± s.e.m.; lines and bands, linear fit and 95% CI. The identity line applies only to calibrated age predictions in A. Scores in B are not calibrated ages. *n* is given in each panel.

We then applied the fixed weights of our proteomic clocks to the same profiles without retraining. Between 91% and 98% of the core protein-clock genes were detected in each organ. Protein-weighted transcriptomic scores increased with age in liver (ρ = 0.90), kidney (0.81), spleen (0.68), brain levels 1 and 4 (0.79 and 0.83) and the pooled brain profile (0.62), and each of these associations passed both the age-permutation test (P ≤ 0.039) and the expression-matched null (P_null ≤ 0.035; ***Fig. 5B***). Brain level 2 showed a similar correlation (ρ = 0.73; P = 0.015) but did not clearly outperform matched random genes (P_null = 0.056), and levels 3 and 5 were marginal on both tests (ρ = 0.61 and 0.57; P = 0.051 and 0.069). In heart and lung, the protein-derived scores were weak (ρ = 0.19 and 0.40; P = 0.56 and 0.27), even though RNA clocks trained on the same data predicted age well (r = 0.91 and 0.79; ***Fig. 5A***). Thus, these spatial profiles contained age-associated RNA information, but the fixed protein-derived weights did not recover it reliably in heart or lung. For heart, this agrees with the bulk analysis, in which protein scores tracked age more strongly than RNA scores (Fig. 4C). Lung, in contrast, transferred well in bulk RNA (***Fig. 4A***), so its weak spatial result may reflect the small number of lung mice (n = 10) or differences between cohorts.

A section-level score can change because expression shifts within a compartment or because compartments with different scores change in relative abundance. To examine these possibilities, spatial spots were grouped into transcriptionally defined niches using 2,000 highly variable genes. Counts were aggregated within each niche for each mouse and scored using the fixed proteomic weights. Age associations were then assessed separately for niche scores and relative niche abundance. Niche scores were positively associated with age in all 65 niches, and the association reached P < 0.05 in 28 niches from brain, kidney, liver and spleen (***Supplementary Fig. 6A***). In 26 of these 28 niches, the share of the section occupied by the niche did not change detectably with age. Across all 65 niches, the age trend in score was unrelated to the age trend in share (ρ = -0.03, P = 0.79**; *Supplementary Fig. 6B***).

**Fig 6.**
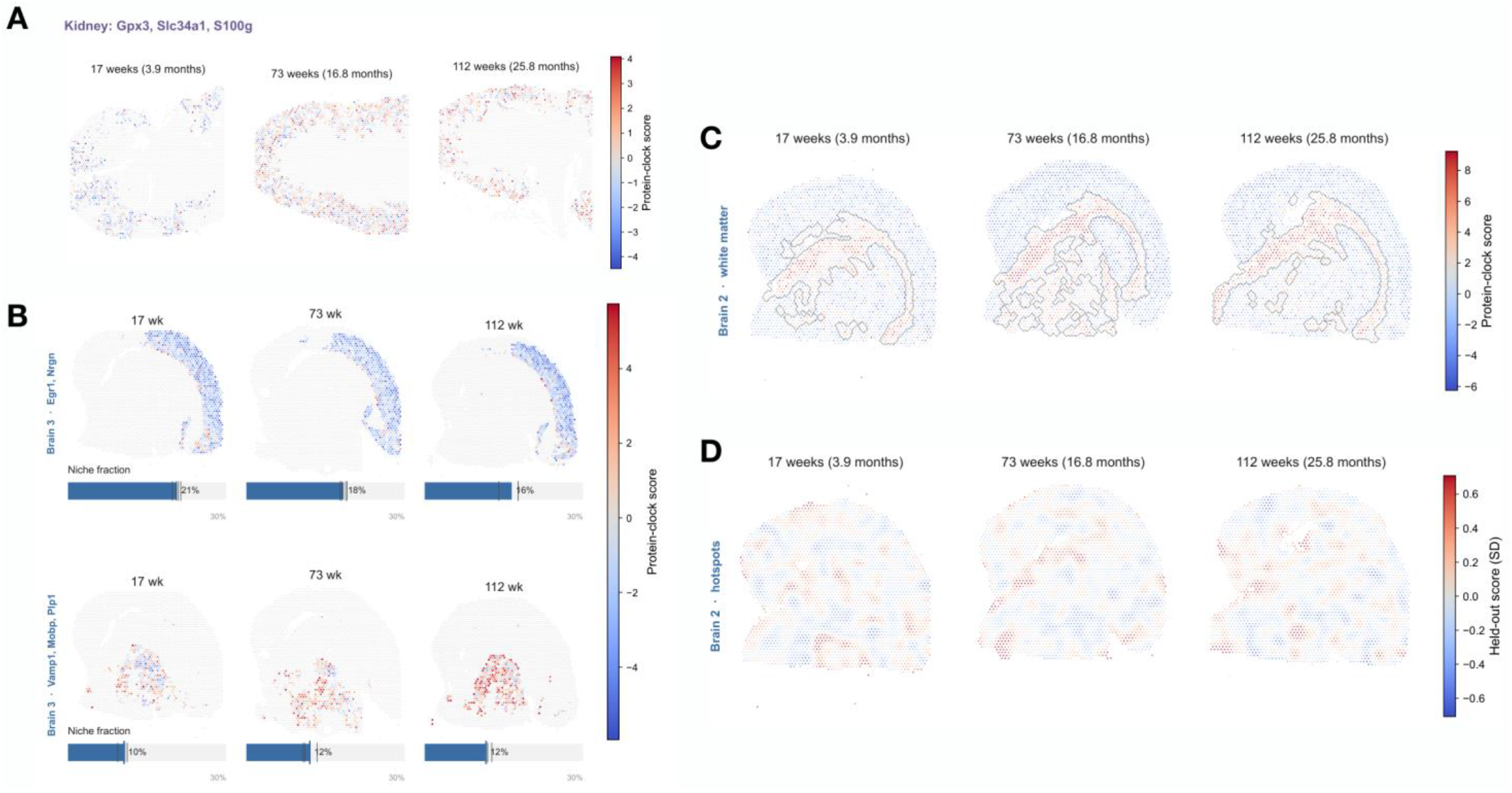
Proteomic aging signatures localize to defined tissue niches and brain microdomains. Representative Visium sections, one mouse per age (Kern et al.^19^). **(A)** Kidney tubular niche (*Gpx3, Slc34a1, S100g*). **(B)** Brain level 3 niches enriched for *Egr1* and *Nrgn* (top) or *Vamp1, Mobp* and *Plp1* (white matter; bottom); bars, niche fraction of the section. In A and B, niche spots are colored by protein-derived RNA score and other spots are grey. **(C)** Protein-derived RNA scores at brain level 2; outlines, white-matter niches. Scores in A–C use RNA expression standardized across spots within each section; the maps show relative spatial localization and do not provide a common calibrated age scale across sections. **(D)** Hotspot map at brain level 2, scored with held-out clock genes in s.d. units (Methods). Color scales differ between panels. Inferential analyses use mouse-level summaries, not individual spots as independent replicates.

Representative niches illustrate this pattern. In kidney, a tubular niche marked by *Gpx3, Slc34a1* and *S100g* showed a strong age-associated increase in its proteomic clock-derived score (ρ = 0.90, P = 0.001), despite no detectable change in its relative abundance (ρ = 0.04; ***Fig. 6A***). Similarly, a periportal liver niche marked by *Cyp2f2, Hsd17b13* and *Sds* exhibited an increase in score with age (ρ = 0.93), without a statistically significant change in its proportion of the section (ρ = ™0.54, P = 0.12; ***Supplementary Fig. 6A***). In the brain, white-matter niches showed the same pattern at three levels (score ρ = 0.80–0.84; share |ρ| ≤ 0.05). At level 3, for example, a white-matter niche marked by *Vamp1, Mobp* and *Plp1* occupied about 10–12% of the section at all three ages while its score rose [ρ = 0.84; P = 0.002] (***Fig. 6B***, bottom). The clearest exception was an *Egr1*- and *Nrgn*-enriched niche, also at level 3, whose score rose with age (ρ = 0.62, P = 0.048) while its share of the section fell from about 21% to 16% (ρ = ™0.94, P < 0.001; ***Fig. 6B***, top). This niche showed increasing clock scores despite a declining share of the section. Together, these results localize age-associated increases in the RNA component of the proteomic signatures to persistent tissue niches, with little accompanying change in their relative abundance.

### 2.5 White matter and glial microdomains exhibit elevated proteomic aging signatures

Finally, we asked whether particular regions had consistently higher scores than their surroundings. We searched every organ but found such regions only in the brain, where they occurred at two scales. In heart, kidney, liver, lung and spleen, no localized hotspots were detected (score excess relative to randomly selected spots within the same niche, ™0.05 to 0.06 s.d.; all P ≥ 0.31). At the broad scale, white-matter niches, defined by *Plp1, Mbp* or *Mobp* among their marker genes, scored higher than the rest of the section in all 55 brain sections (***Fig. 6C***). The difference averaged 3.8–6.8 score units across brain levels and age groups and exceeded 3.0 units in every section. It did not change significantly with age at any brain level (P = 0.077–0.55), so it was already present at 3.9 months.

Discrete hotspots were also detected within brain niches. Hotspots were identified using one randomly selected half of the clock genes and evaluated using the held-out half. Each brain section contained an average of 9–15 candidate hotspots; when evaluated using the held-out genes, these spots still scored 0.18 s.d. higher than randomly selected control spots within the same niche (brain levels averaged within each mouse, P < 0.001; levels 1–4, P ≤ 0.039; level 5, P = 0.075; ***Fig. 6D***). Tissue within 150 µm of these hotspots showed higher expression of reactive astrocyte-associated genes (*Vim, C4b, Aqp4* and *Padi2*) and of microglial and MHC class I-associated genes (*Lyz2, C1qa, H2-K1* and *B2m*), and lower expression of neuronal markers (*Kcnh3* and *Rbfox3*). Within individual brain niches, the proteomic signature therefore concentrates at discrete, glia-enriched, and neuron-poor microdomains. Hotspot enrichment persisted after adjustment for local oligodendrocyte, neuronal and astrocyte marker expression (score excess, 0.10 s.d.; P < 0.001; levels 1–4, P ≤ 0.023).

We then asked whether these hotspots changed with age. They were already detectable at 3.9 months, and their number per section did not change significantly with age at any brain level (P ≥ 0.18). After adjustment for local oligodendrocyte, neuronal and astrocyte marker expression, hotspot enrichment was not associated with age overall (ρ = 0.18, P = 0.60). A nominal increase at level 2 (ρ = 0.62, P = 0.041) did not survive correction across the five levels, and without adjustment the overall association was also only nominal (ρ = 0.59, P = 0.049), with inconsistent trends across levels. These findings distinguish age-associated increases within tissue niches from regional enrichment of the protein-derived aging signature already present in young adulthood.

## 3. Discussion

Proteomic aging signatures differ across organs in their molecular composition and their representation in RNA. Clocks developed from eight mouse organs generalized across independent proteomic cohorts, and their fixed protein-derived weights recovered age-associated patterns in bulk and spatial transcriptomes. Spatial analysis further distinguished age-associated increases within defined niches from persistent enrichment in white matter and glia-associated microdomains.

Previous mouse atlases have identified shared alterations in immunity and metabolism alongside substantial differences among organs^12,13^. In our models, 93% of core clock proteins were selected in only one organ, and most signatures transferred poorly between organs even when analysis was restricted to proteins detected in all eight tissues. This combination supports organ-dependent aging patterns beyond differences in protein coverage. However, selection among correlated proteins can produce different predictors for related biological processes, so limited feature overlap should not be equated with entirely separate aging mechanisms. The inclusion of proteins without individually significant age associations also shows that the clocks capture combined patterns that extend beyond univariate differential-expression testing. Their selection establishes predictive contributions rather than a causal role in aging. Human plasma studies similarly associate organ-enriched protein signatures with distinct health outcomes^1^, supporting the value of resolving aging at the organ level.

Applying the same selected genes and coefficients across molecular layers tests whether an aging pattern learned from proteins is also represented in RNA abundance. Reproducible transfer across independent transcriptomic cohorts indicates that an age-associated component of the proteomic signatures is shared with the transcriptome. The gene-set controls strengthen this interpretation, particularly in kidney, liver and lung, where the selected genes outperformed matched alternatives. Moreover, retaining only the direction of the protein-derived weights largely preserved age associations, whereas assigning every gene an equal positive contribution weakened them. Thus, transfer depended mainly on the selected genes and their signed contributions rather than the precise coefficient magnitudes. The small overlap between independently trained spatial RNA clocks and proteomic clocks further suggests that different predictor sets can recover related age-associated information.

Protein scores were more strongly associated with age than matched RNA scores in brain and skeletal muscle. Heart showed the same direction of difference at nominal significance. Because the paired analyses used matched mice, genes and weights, these differences cannot be attributed to independent clock training or differences between cohorts. They accord with reports of age-associated transcriptome–proteome divergence in mouse and human tissues, including changes in extracellular matrix and mitochondrial proteins that are incompletely reflected in RNA abundance^2,11^. Long-lived mitochondrial proteins provide one plausible mechanism through which protein abundance and function can become temporally separated from transcript abundance^23^. Age-dependent changes in translation and degradation are additional candidates, although their contributions were not measured here. Conversely, comparable age associations in kidney and liver indicate substantial RNA representation of these particular protein signatures.

Spatial transfer localized the RNA component of proteomic aging signatures to defined tissue niches. Single-cell aging atlases and spatial transcriptomic clocks have shown that age-associated expression changes depend on cell identity and local cellular environments^16,18^. Here, the signatures were defined independently in tissue proteomes and applied to spatial RNA without retraining. Scores increased with age within kidney tubular, hepatic and white-matter niches without detectable changes in their relative representation. These findings support molecular remodeling within established tissue compartments as a contributor to whole-organ aging signatures.

White matter and localized glia-associated microdomains exhibited elevated scores relative to their surroundings. White-matter enrichment was already present in young adults, distinguishing this regional offset from the age-associated increases observed within niches. Hotspots were enriched for reactive astrocyte, microglial and immune-associated transcripts, consistent with processes implicated in brain aging, including astrocyte reactivity and microglial handling of myelin debris^24,25^. Hotspot score enrichment was confirmed using held-out clock genes and persisted after adjustment for niche identity and local oligodendrocyte, neuronal and astrocyte marker expression, supporting localized molecular variation beyond broad compositional differences. Higher regional scores alone do not establish older biological age or faster local aging.

Rapamycin was associated with lower clock scores in male skeletal muscle and female liver in the Burns cohort, and in male liver in the Watanabe cohort. Rapamycin extends lifespan in genetically heterogeneous mice^26^, and previous work has shown that liver DNA methylation aging signatures respond to rapamycin and other lifespan-extending conditions^27^. Our findings extend these observations to selected tissue proteomic signatures and reveal differences in molecular responses across organs and cohorts. The GHRKO and Snell dwarf comparisons produced mixed responses, indicating that intervention responsiveness requires evaluation alongside chronological-age prediction.

Human plasma proteomic clocks have been associated with incident disease and mortality, illustrating the potential biological relevance of proteomic age measures^9^. Establishing comparable relevance for the present tissue signatures will require prospective functional validation. The external scores were standardized within each dataset and quantify relative age-associated patterns rather than calibrated age in months, limiting comparisons of absolute values across cohorts or molecular layers. Spatial scores represent the RNA component of the protein-derived signature, and Visium resolution leaves finer changes in cellular composition unresolved. Most analyses used male mice on C57BL/6 backgrounds, so extension to females and diverse genetic backgrounds remains an important next step. The spatial cohort also contained only three age groups, restricting resolution of nonlinear trajectories and differences in timing between niches. Larger cohorts with paired RNA and protein measurements would help determine whether the observed organ differences persist across sex, genetic background and later-life pathology.

Proteomic aging signatures are predominantly organ-specific and differ in their correspondence with RNA. Their projection onto spatial transcriptomes distinguishes age-associated remodeling within tissue niches from persistent regional enrichment, connecting whole-organ proteomic aging signatures to their anatomical context.

## 4. Methods

### 4.1 Proteomic datasets and preprocessing

Discovery proteomes were obtained from Scifo et al.^12^ and comprised brain, heart, lung, liver, kidney, spleen, gastrocnemius muscle and testis from male C57BL/6J mice aged 3, 5, 8, 14, 20 and 26 months (44 mice per organ after preprocessing). Contaminants and non-mouse proteins were excluded. Proteins detected in at least 80% of mice within an organ were retained, log2-transformed and median-centred within samples. Duplicate accessions were represented by the first record. Independent aging proteomes were obtained from the tandem mass tag dataset of Takasugi et al.^11^ and the PulseDIA dataset of Lu et al.^14^, using male mice aged 6–30 and 1–24 months, respectively. Published quantification matrices were retained, protein identifiers were reduced to primary accessions, and duplicate accessions were collapsed by the median. Takasugi proteins with a reported false-discovery rate ≥0.05 were excluded. Hippocampus and gastrocnemius were matched to the brain and muscle clocks, respectively. Longevity-intervention analyses used gastrocnemius, liver and kidney proteomes from male and female UM-HET3 mice receiving rapamycin, acarbose, canagliflozin, 17α-estradiol or calorie restriction, together with growth hormone receptor knockout (GHRKO) and Snell dwarf comparisons from Burns et al.^20^. The GHRKO analysis used male C57BL/6 mice. Liver rapamycin data were also obtained from the Watanabe et al. LC-M001 series^21^, comparing 14-ppm rapamycin with the 5LG6 control diet. Repeat aliquots were excluded, protein intensities were summarized by primary accession using the median, and positive values were log2-transformed; non-positive values were treated as missing. Treatment comparisons were performed within organ and sex, and each genetic model was compared with its corresponding wild-type controls. Watanabe males and females were standardized separately.

### 4.2 Clock development and validation

Separate elastic-net age predictors were fitted for each organ using scikit-learn ElasticNetCV. Within each training fold, missing values were imputed by the protein median and features were standardized to zero mean and unit variance. Penalties were selected by five-fold cross-validation over mixing parameters of 0.2, 0.5 and 0.8 and ten logarithmically spaced penalty strengths from 10^−2^ to 10^1^ (tolerance, 10^−3^; maximum iterations, 8,000). Performance was evaluated by leave-one-mouse-out cross-validation, repeating imputation, scaling and penalty selection without the held-out mouse. Prediction accuracy was summarized by Pearson correlation and mean absolute error. Core clock proteins were those with non-zero coefficients in more than half of the leave-one-mouse-out folds. Their fixed weights were the mean non-zero coefficients across folds. Protein contributions were summarized as the mean absolute product of standardized abundance and fixed weight across mice. For cross-organ proteomic prediction, models were restricted to proteins passing the 80% detection filter in all eight organs. Missing values were imputed using organ-specific protein medians calculated across all mice before cross-validation. Models trained in one organ were applied to the corresponding target-organ sample from the held-out mouse. Correlations with chronological age were calculated for within-organ and cross-organ predictions.

### 4.3 Transfer of proteomic signatures to independent datasets

Independent proteomic and RNA datasets were scored using detected core clock features and their fixed weights, without retraining. Missing entries were imputed by the median of observed clock-feature abundances within the same sample. Features were standardized across samples within each organ and analysis cohort to zero mean and unit population standard deviation; zero-variance features were excluded. Scores were calculated as the weighted sum of standardized abundances, requiring at least three detected features. These scores quantify relative molecular signatures and were not converted to age in months. Bulk transcriptomes were obtained from Takasugi et al.^11^ (GSE225576) and Shavlakadze et al.^22^ (GSE314817). Takasugi RNA matrices were DESeq2-normalized counts, and Shavlakadze RNA matrices were ArrayStudio gene counts. Both were transformed to log2(x + 1). Duplicate gene symbols in the Shavlakadze matrix were collapsed by the median before this transformation. Takasugi gene identifiers were already unique, and every value in both matrices was at least zero. Protein accessions were mapped to annotated gene symbols, and weights mapping to the same gene were summed. Cortex was scored with the brain clock and left ventricle with the heart clock. Cross-organ RNA transfer applied each weight vector separately to each target tissue, using genes shared by the weight vector and that target matrix. For paired protein–RNA comparisons, Takasugi samples were matched by age and replicate, and both scores used identical mice, genes and weights; protein abundances mapping to one gene were summarized by the median.

### 4.4 Differential abundance and signature controls

Age-associated protein abundance was assessed in the discovery matrices using a moderated F test with age as a six-level factor. Missing values were imputed by the protein mean, and residual variances were moderated towards the median variance with four prior degrees of freedom. *P* values were adjusted within each organ using the Benjamini–Hochberg procedure, with *q* < 0.05 defining age-differential proteins. These proteins were compared with the core clock sets. RNA signature specificity was assessed against 1,000 non-clock gene sets of equal size sampled randomly or matched without replacement for mean expression or expression variance. Each set received the same weight vector as the corresponding clock and underwent identical scoring. Spatial RNA controls used expression-matched sets drawn from genes detected in more than half of the mice. Weighting controls applied the exact coefficients, coefficient signs alone, or equal positive weights to the same clock genes.

### 4.5 Spatial transcriptomic profiling

Visium data were obtained from Kern et al.^19^ (GSE283283), comprising 111 sections from 15 male C57BL/6JN mice aged 17, 73 and 112 weeks (3.9, 16.8 and 25.8 months). Tissues included brain, heart, kidney, liver, lung and spleen; brain was sampled at five coronal levels (bregma −1.7, −2.06, −2.7, −3.52 and −4.72 mm). Space Ranger filtered matrices were used, and counts sharing a gene symbol were summed. Counts were aggregated across section spots to obtain one profile per mouse and transformed to log2(CPM + 1), where CPM denotes counts per million. Pooled brain profiles were generated by centring each level by its gene means and averaging available levels within each mouse. RNA clocks used the 5,000 most variable genes among those with ≥1 CPM in at least 80% of sections; pooled brain eligibility used detection fractions weighted by section number. Models used the elastic-net specification and validation described above. Additional models trained on young and old mice were evaluated in held-out middle-aged mice. RNA genes selected in more than half of the validation folds were compared with matched proteomic core genes among the 5,000 genes tested. Proteomic signatures were transferred to section and pooled brain profiles using the fixed scoring procedure. Gene symbols were matched without regard to case.

### 4.6 Tissue niche analysis

Transcriptional niches were identified separately for each organ or brain level using the 2,000 most variable genes after log(1 + counts per 10,000) transformation. Variance was calculated as the spot-weighted mean of within-section variances. Expression was centred within sections, and the leading 30 principal components were clustered using mini-batch k-means with ten initializations and ten clusters per brain level or eight per other organ. Niches were annotated by the three genes with the largest mean expression differences between niche and non-niche spots. Niche counts were aggregated within each mouse, requiring at least 30 spots per profile and eligible profiles from at least 80% of mice, rounded up. Profiles were transformed to log2(CPM + 1) and scored with the corresponding fixed proteomic weights. Niche abundance was the fraction of section spots assigned to that niche, including zero fractions when absent. Score and abundance associations with age were evaluated separately, and their correlations across niches were compared by Spearman correlation with permutation testing. Niches marked by *Gm42418, Lars2* or *Malat1* were classified as technical and excluded from biological summaries.

### 4.7 Spatial enrichment and hotspot analysis

Spot scores were calculated from log2(CPM + 1) expression standardized across spots within each section, using at least three variable clock genes. White-matter niches were identified by *Plp1, Mbp* or *Mobp* markers, and enrichment was calculated as the mean score difference between white-matter and remaining spots.

Hotspots were identified after regressing out niche identity and log library size and scaling residual scores to unit standard deviation. Additional brain analyses adjusted for mean log(1 + counts per 10,000) expression of oligodendrocyte markers (*Plp1, Mbp, Mobp, Mag, Cnp, Cldn11, Mog, Ermn*), neuronal markers (*Snap25, Syt1, Rbfox3, Stmn2, Nrgn*) and astrocyte markers (*Aqp4, Slc1a2, Gja1, Aldoc, Slc1a3*), excluding clock genes. Coordinates were scaled to a Visium spot diameter of 55 µm. Clock genes were randomly divided once into two halves for independent hotspot detection and evaluation. Detection scores were Gaussian-smoothed (standard deviation, 100 µm; truncation, 300 µm). Candidate peaks were local maxima within 300 µm at or above the 95th percentile and had at least 24 neighboring spots within 300 µm. Held-out, unsmoothed scores quantified enrichment as the mean of the peak and 50–150-µm ring minus the mean of the 350–450- and 450–600-µm rings, corrected by the same contrast from 50 sets of random centers matched for niche and neighboring spot count. Gene-half assignments were reversed and the two estimates averaged; pooled brain estimates averaged available levels within each mouse. Hotspot-associated expression was measured as log2 fold change between CPM-normalized counts aggregated within 150 µm of peaks and matched control centers. Clock genes and genes below 20 CPM in matched controls were excluded.

### 4.8 Statistical analysis

Mice were the biological replicates for cohort comparisons. Associations of transferred scores and niche fractions with age were tested using two-sided Spearman permutation tests with 10,000 age-label permutations; white-matter and hotspot age associations used 5,000 permutations. RNA clock performance used 200 age-label permutations with model refitting and a one-sided comparison of out-of-fold Spearman correlations. Gene-set null comparisons were also one-sided. Permutation *P* values used the add-one correction, *P* = (*b* + 1)/(*B* + 1), where *b* is the number of equally or more extreme permutations among *B* permutations. Intervention effects were assessed using two-sided permutation tests of the Mann–Whitney U statistic (10,000 group-label permutations) and Cliff’s δ, with negative values indicating lower treated scores. Paired protein and RNA Spearman correlations with age were compared using two-sided Steiger tests accounting for their overlapping correlation. Mean hotspot enrichment was assessed by exact sign-flip tests of mouse-level estimates. Intervention and Steiger *P* values were unadjusted. Analyses used Python with pandas, NumPy, SciPy, h5py and scikit-learn, with fixed random seeds.

## Data availability

Proteomic data analyzed in this study can be downloaded from ProteomeXchange, PRIDE and jPOST, and transcriptomic data from the Gene Expression Omnibus. Accession numbers for each dataset are listed in Supplementary Table 1. Source data are provided with this paper.

## Code availability

Code used in this study is available at: github.com/sean6211/mice-organ-specific-proteomic-aging-clocks.

## Acknowledgements

This work was supported by the National Institute on Aging of the National Institutes of Health (R01AG065301, R21AG087381, R21AG094596, R21AG099390, K07AG097560 and R41AG099556 to A.M.P.). We thank the investigators who generated and publicly shared the proteomic and transcriptomic datasets analyzed in this study.

