## Supplemental Tables and Figures for "Proteomic clocks reveal organ-specific signatures of aging in mice"

Public accessions of the datasets analyzed in this study.

| Dataset | Repository | Accession |
|---|---|---|
| Scifo et al. discovery proteomes | ProteomeXchange | PXD058684 |
| Scifo et al. discovery proteomes | jPOST | JPST003472 |
| Takasugi et al. proteomes | PRIDE | PXD040722 |
| Takasugi et al. RNA-seq | Gene Expression Omnibus | GSE225576 |
| Lu et al. proteomes | PRIDE | PXD053154 |
| Burns et al. proteomes | PRIDE | PXD040497 |
| Watanabe et al. liver proteomes | PRIDE | PXD035255 |
| Shavlakadze et al. bulk RNA-seq | Gene Expression Omnibus | GSE314817 |
| Kern et al. Visium | Gene Expression Omnibus | GSE283283 |

**Supplementary Table 1.** Public accessions of the datasets analyzed in this study.

### Supplementary Figures

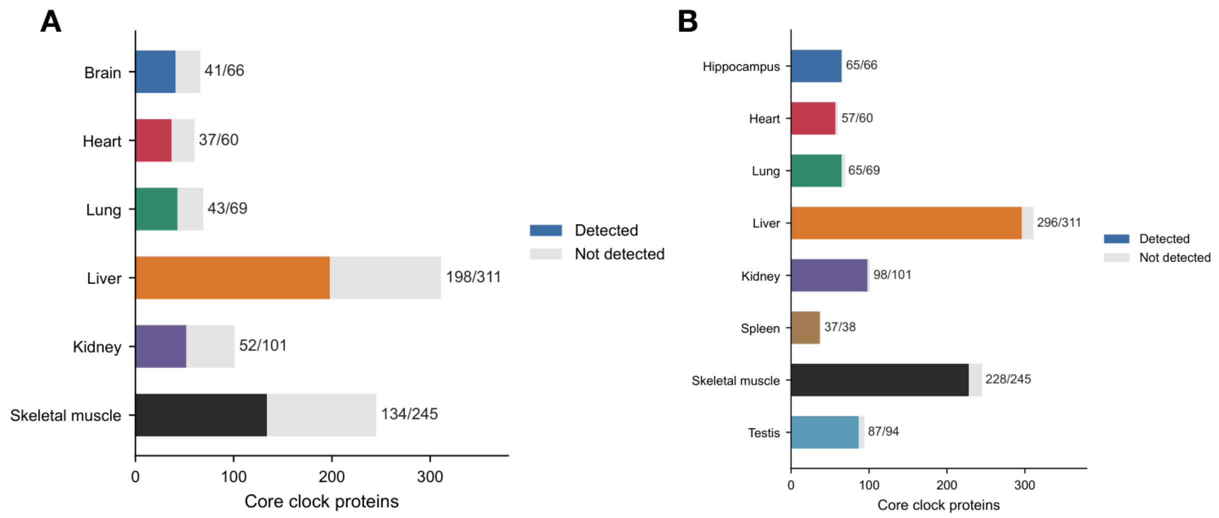


**Supplementary Figure 1. Coverage of core clock proteins in independent proteomes.** Numbers of core clock proteins detected (organ colours) or not detected (grey) in the Takasugi et al.[11] **(A)** and Lu et al.[14] **(B)** datasets. Labels give detected proteins relative to the total number in each core clock. Core proteins were selected in more than half of the discovery leave-one-mouse-out folds. The brain clock was matched to hippocampus in the Lu dataset.

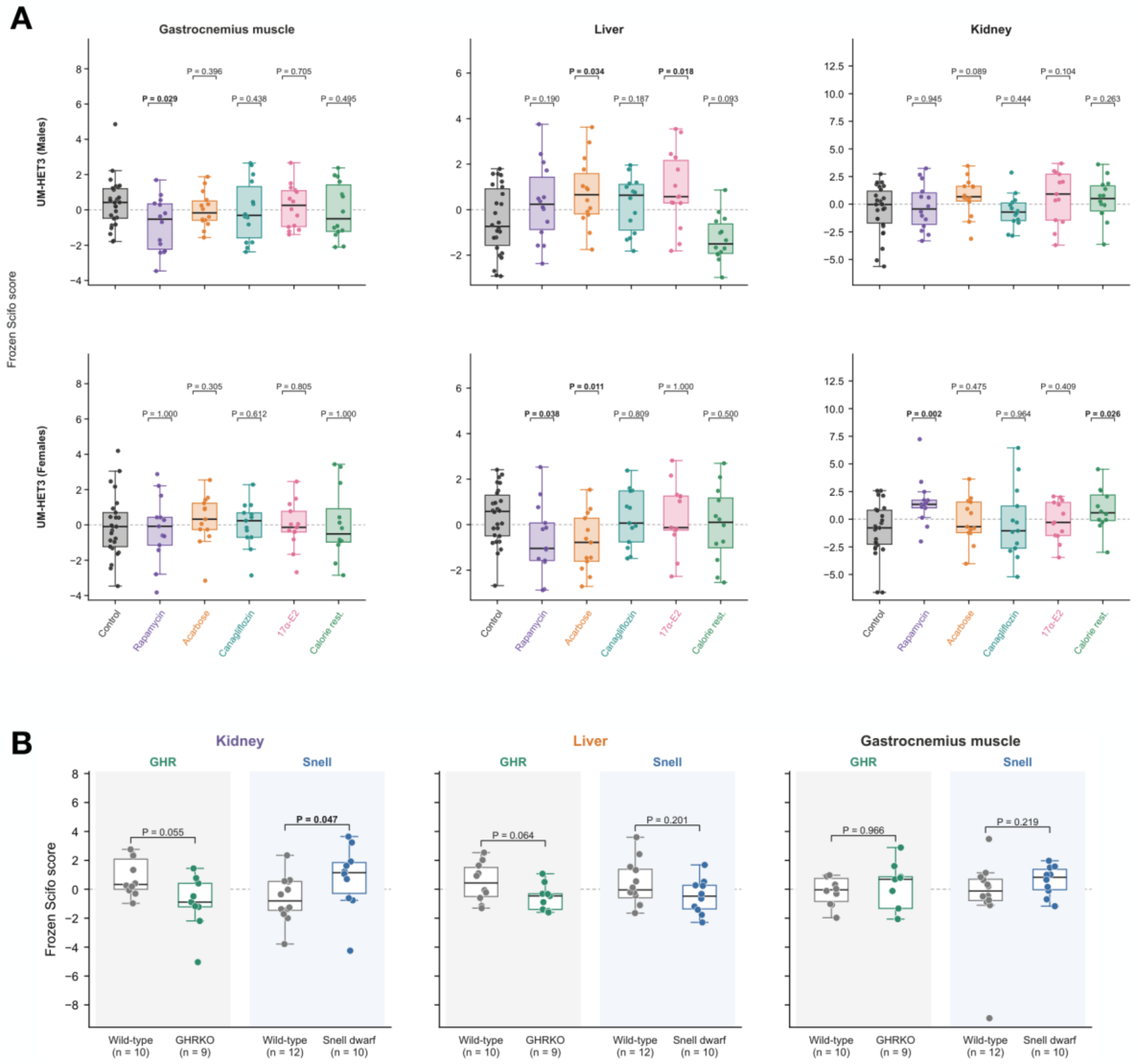


**Supplementary Figure 2. Organ clock scores under pharmacological, dietary and genetic longevity interventions.** **(A)** Fixed clock scores in gastrocnemius muscle, liver and kidney from 12-month-old UM-HET3 mice in the Burns et al.[20] dataset. Males are shown above and females below. Groups are control, rapamycin, acarbose, canagliflozin, 17α-estradiol and calorie restriction. Male groups contain 26 controls and 14 mice per intervention, except for the kidney 17α-estradiol group (*n* = 13). Female controls number 25 for muscle and 26 for liver and kidney; female intervention groups contain 13 mice, except calorie restriction (*n* = 12). Each *P* value compares the indicated intervention with the same-sex, same-organ control group. **(B)** Kidney, liver and gastrocnemius scores in growth hormone receptor knockout (GHRKO) mice and Snell dwarf mice, with their respective wild-type controls. GHRKO comparisons contain 10 wild-type and 9 knockout mice; Snell comparisons contain 12 wild-type and 10 dwarf mice per organ. The GHRKO analysis used male mice. Points represent individual mice; boxes show the interquartile range with the median marked. Scores use fixed proteomic weights and are not calibrated ages. *P* values are from two-sided permutation tests of the Mann–Whitney U statistic (10,000 group-label permutations); bold values indicate unadjusted $P < 0.05$.

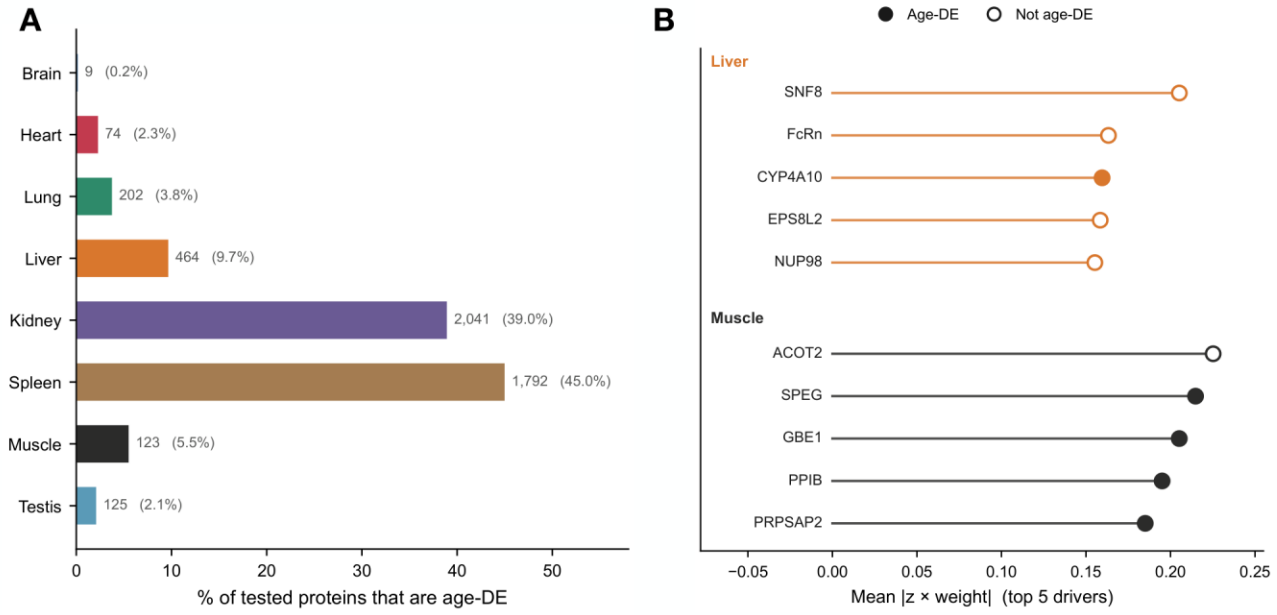


**Supplementary Figure 3. Core clock proteins include predictors without significant individual age associations**. Analyses used the discovery proteomes of Scifo et al.[12]**. (A)** Number and percentage of tested proteins that were age-differential in each discovery organ ($n$ = 44 mice per organ). Age was modelled as a six-level factor using a moderated F test, with Benjamini–Hochberg $q < 0.05$ within each organ. **(B)** The five core clock proteins with the largest contributions in liver and skeletal muscle, ranked by the mean absolute product of standardized protein abundance and fixed clock weight across mice. Filled circles denote age-differential proteins; open circles denote proteins not meeting the $q < 0.05$ threshold.

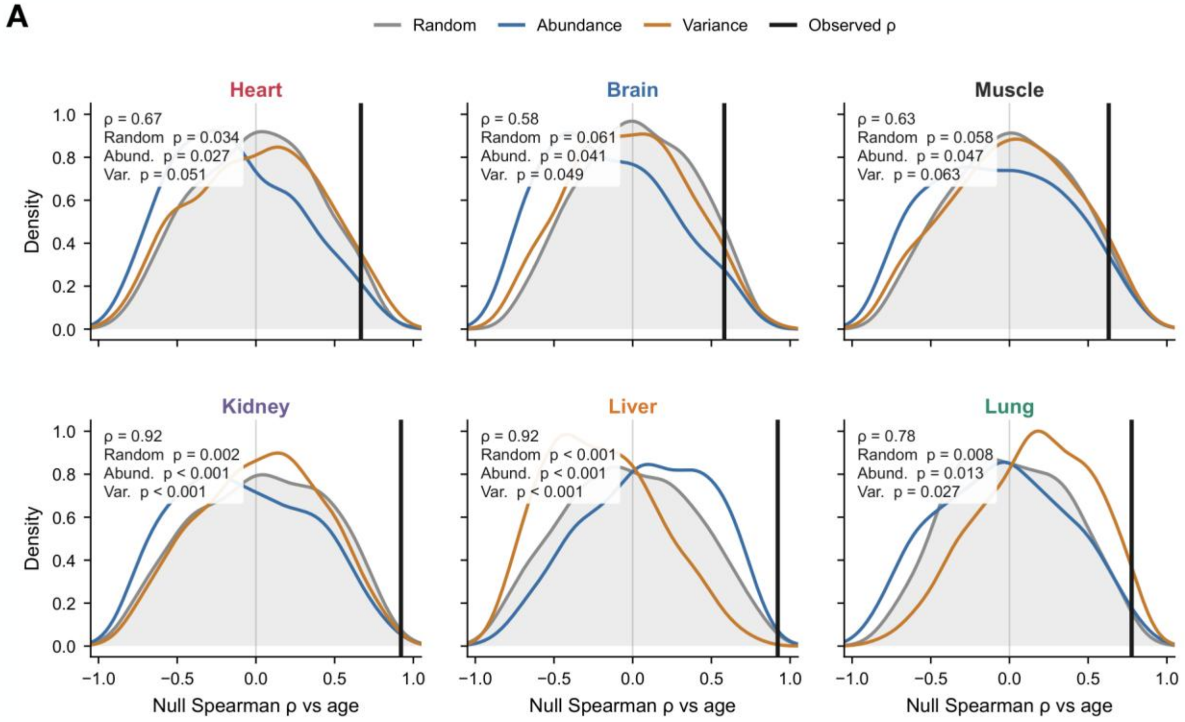


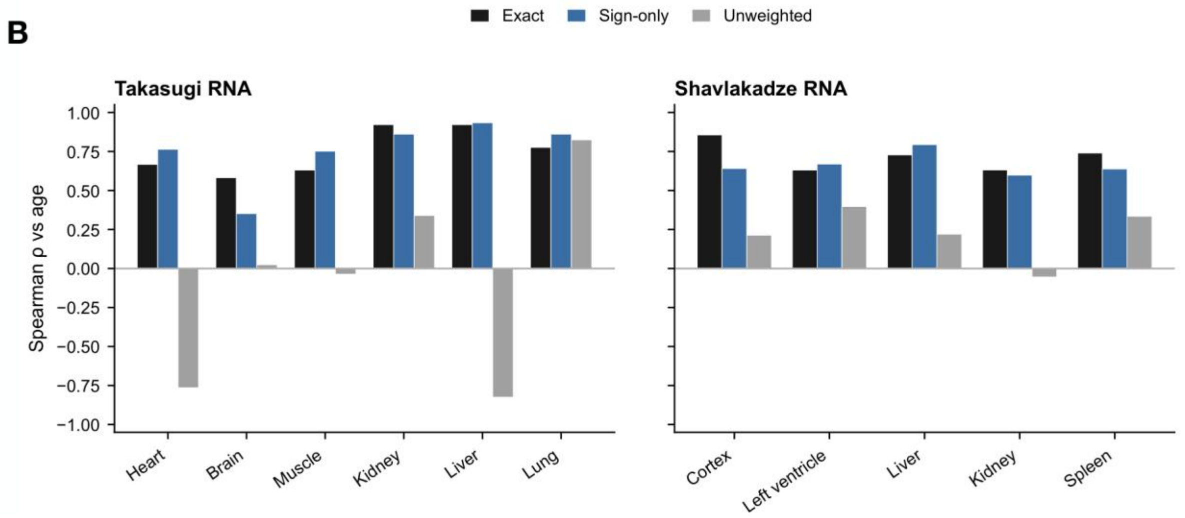


**Supplementary Figure 4. Gene selection and coefficient direction contribute to transfer of proteomic signatures to RNA. (A)** Spearman correlations between RNA clock scores and age in the Takasugi et al.[11] cohort (black vertical lines; *n* = 16 mice per organ), compared with null distributions from 1,000 gene sets sampled randomly (grey), matched for mean expression (blue), or matched for expression variance (orange). Each null set contained the same number of genes as the corresponding detected clock-gene set and carried the same weight vector. *P* values are one-sided comparisons with the corresponding null distribution. **(B)** Spearman correlations with age for RNA scores using the exact fixed coefficients (black), coefficient signs only (blue), or equal positive weights (grey), applied to the same clock genes in the Takasugi (left) and Shavlakadze et al.[22] (right) cohorts. Cortex uses the brain clock and left ventricle the heart clock. Scores were computed without retraining.

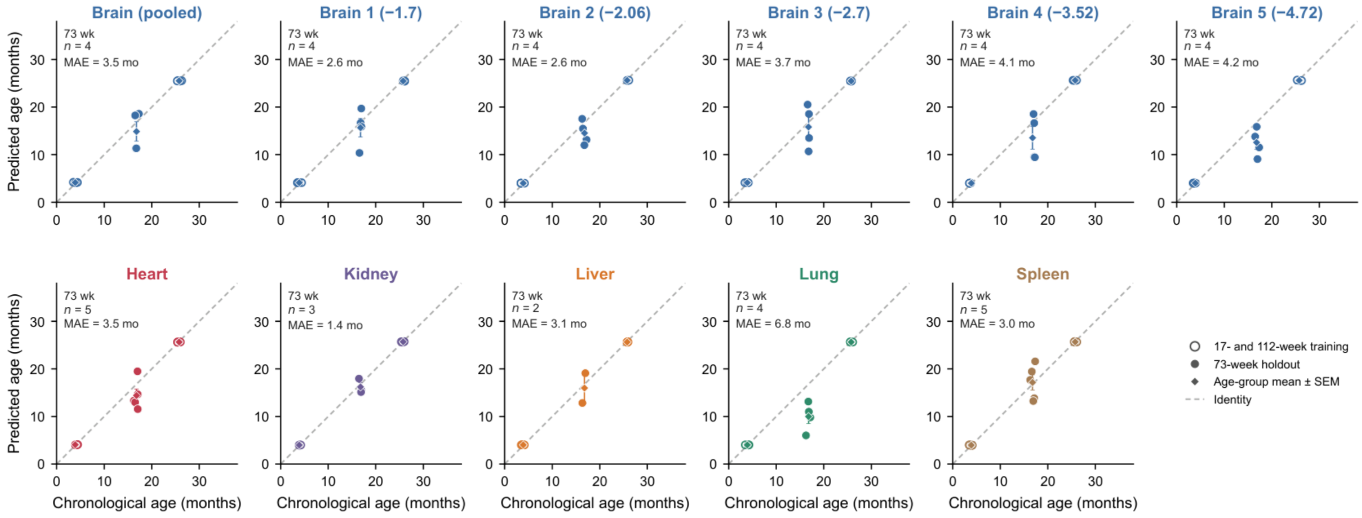


**Supplementary Figure 5. Prediction of held-out middle-aged mice by spatial RNA clocks.** Elastic-net RNA clocks were trained on summed Visium profiles from 3.9- and 25.8-month-old mice (open circles) and applied to held-out 16.8-month-old mice (filled circles). Visium data were obtained from Kern et al.[19], with profile construction as in Fig. 5A. Panels report the number of held-out mice and their MAE in months. Held-out sample sizes are four for pooled brain and each brain level, five for heart, three for kidney, two for liver, four for lung and five for spleen. Diamonds and error bars show age-group mean ± s.e.m.; dashed lines indicate identity. Brain levels are labelled by bregma coordinate in millimetres. Statistics shown in each panel concern the held-out middle-aged mice.

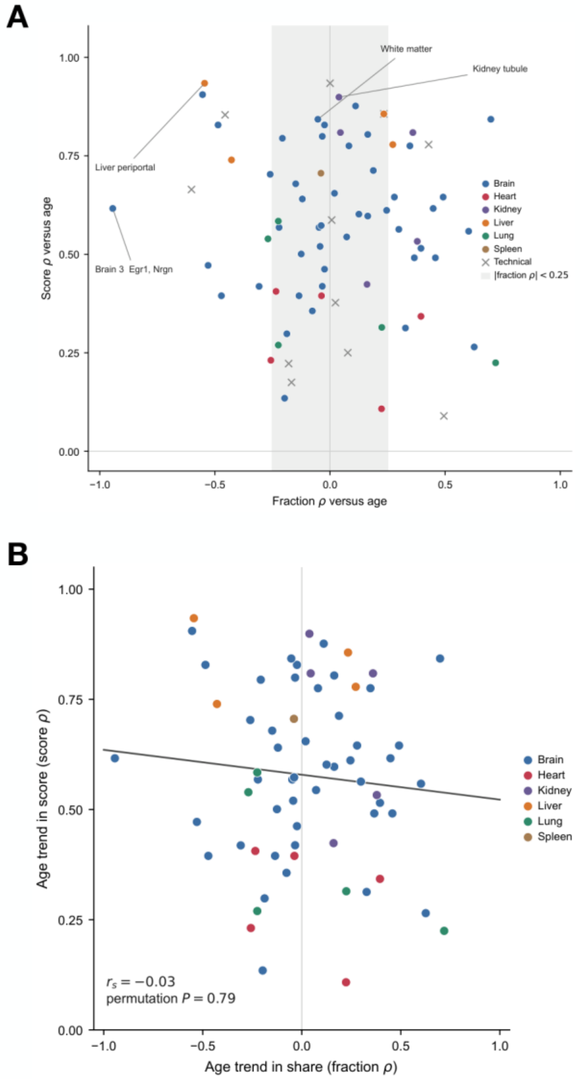


**Supplementary Figure 6. Niche scores increase with age across diverse changes in niche abundance**. Analyses used Visium data from Kern et al.[19]**. (A)** For each transcriptionally defined niche, Spearman $\rho$ between its fraction of section spots and age (horizontal axis) is plotted against Spearman $\rho$ between its protein-derived clock score and age (vertical axis). Niche scores were obtained by summing RNA counts across niche spots within each mouse and applying the fixed organ-specific proteomic weights without retraining. Only niches with at least 30 spots in at least 80% of mice were eligible for scoring. Colours identify organs; grey crosses identify technical niches whose marker genes included *Gm42418*, *Lars2* or *Malat1*. The shaded band marks $|\text{fraction } \rho| < 0.25$. Labelled examples identify white matter, kidney tubule, periportal liver and the *Egr1*/*Nrgn*-enriched brain level 3 niche. (**B)** Association between the two age correlations across the 65 non-technical niches. Each point represents one niche; the solid line is a descriptive linear fit. The indicated Spearman correlation and permutation *P* value assess the association across niches. Each underlying niche–age correlation was calculated from mouse-level profiles; spots were not treated as independent biological replicates.
